# Buying time – increasing yield potential in wheat by extending stem elongation duration

**DOI:** 10.1101/2025.04.28.650957

**Authors:** Lukas Kronenberg, Oscar E. Gonzalez-Navarro, Sarah Collier, Monika Chhetry, Phillip Tailby, Michelle Leverington-Waite, Luzie U. Wingen, Simon Griffiths

## Abstract

Extending the duration of the stem elongation (SE) phase between terminal spikelet (TS) and anthesis has often been proposed as an avenue to increase wheat yield. However, accurate determination of TS is labour intensive, and existing evidence is often based on a limited number of genotypes observed under controlled conditions.

Here, a Buster x Charger population comprising 108 doubled haploid lines was grown across four year-sites under UK field conditions. TS was recorded through meristem dissection and SE duration was measured as the time between TS and ear emergence (EE). Mixed model analysis across year-sites revealed high heritabilities (H^2^_yield_ = 0.66, H^2^_TS_ = 0.93, H^2^_SE_ = 0.89, H^2^_EE_ = 0.95) and strong genetic correlations between yield and the phenology traits SE and EE (*r*_*g*_ = 0.56 and *r*_*g*_ = 0.6, respectively). While SE duration was mainly driven by EE, independent QTL for TS suggest that SE could be modified without affecting EE. The positive effect of SE duration on yield was attributed to increased grain number per area as well as increased grain weight, through independent QTL. Although validation in broader genetic backgrounds is needed, these QTL may offer opportunities for further yield increase in a physiological breeding framework.

**Highlights:**

- Longer stem elongation duration increases grain number.
- Longer stem elongation duration increases grain weight.
- Independent QTL suggest the timing of stem elongation as well as its effect on yield components may be independently selectable.

## Introduction

The appropriate timing of a crops life cycle, defined by key phenological events such as flowering time, is paramount for its performance and yield potential in a given environment (Snape *et al*., 2001; Langer *et al*., 2014). Due to global warming, bread wheat (*Triticum aestivum*) heading or flowering dates have advanced from 0.8 up to 2.86 days per decade in the past century, depending on the examined locations and time-scales (Hu *et al*., 2005; Tao *et al*., 2006; Wang *et al*., 2015). Furthermore, yield increase stagnation has been reported in wheat as well as other major crop species (Ray *et al*., 2012; Schauberger *et al*., 2018). Further yield increase is critically needed, especially since wheat yields are predicted to decline by 6% for every further degree in temperature increase (Asseng *et al*., 2015).

Grain yield is a complex physiological trait depending on a large number of genetic and environmental factors and their interactions (Reynolds *et al*., 2022; Murchie *et al*., 2023; Slafer *et al*., 2023). Yet, agronomically, it can be broken down into the two numeric components grain number per area and grain weight. Under field conditions, grain number is generally the more important yield determinant (Magrin *et al*., 1993; Slafer *et al*., 2001*a*; Peltonen-Sainio *et al*., 2007). The final number of grains is a function of the components grains per spikelet, spikelets per spike and spikes per plants and is fixed at grain set, just after flowering (Reynolds *et al*., 2022). The number of spikelets per spike is fixed with the formation of the terminal spikelet (TS), which signifies the completion of spikelet initiation, marks the end of the vegetative-to-reproductive transition and concurs with the onset of stem extension (SE) (Slafer *et al*., 2015). Throughout the SE phase, while some of the tillers are aborted, florets are initiated at the spikelets until around booting, when a large proportion of the florets are aborted due to resource competition with the growing stems and spikes towards anthesis (Slafer and Rawson, 1994; Slafer *et al*., 2023). Thus, yield potential is strongly source limited during the SE phase, which leads to a sink limitation of yield during grain filling (Reynolds *et al*., 2022). The introduction of semi-dwarf varieties during the *green revolution* led to a dramatic increase in wheat yields as more resources could be allocated to the spike thanks to the shorter stems, thus increasing harvest index (Slafer *et al*., 2023). Unfortunately, the potential of yield increase through harvest index is approaching its theoretical limit and further progress seems to be difficult (Slafer *et al*., 1999; Reynolds *et al*., 2009).

An often proposed alternative to increase grain number is the extension of the SE duration, to allow for more biomass accumulation and partitioning to the growing spike and lessen competition during SE (Slafer and Rawson, 1994; Slafer *et al*., 1996, 2001*b*, 2015; Miralles and Slafer, 2007; Whitechurch *et al*., 2007; Reynolds *et al*., 2009; Pérez-Gianmarco *et al*., 2018). However, studies investigating the effects of SE duration on yield or yield components were often performed under controlled conditions and evidence from field experiments is mainly based on a limited number of genotypes (see references above, as well as e.g.: Gonzalez-Navarro *et al*., 2016; Guo *et al*., 2018*b,a*; Pérez-Gianmarco *et al*., 2018, 2019; Roychowdhury *et al*., 2023). Data availability from field conditions is limited due to the time and labour expense necessary to phenotype especially the TS stage in field experiments. Proxy measures to determine the timing and growth dynamics of SE have been developed using high throughput phenotyping approaches (Kronenberg *et al*., 2017, 2021; Roth & Kronenberg et al., 2023*b*). However, precise determination of TS as well as a direct relation of the timing of SE to yield is lacking in these studies.

Here, we set out to precisely determine the timing of SE and its relationship to grain yield and yield components in a segregating population comprising 108 doubled haploid lines under field conditions. The aim was to determine whether a prolonged SE duration translates to increased yield and to apply QTL mapping to determine whether an extended SE duration could be achieved by modifying TS or heading independently.

## Materials and Methods

### Plant Material and experimental design

A doubled haploid population of 108 lines was developed from a cross between the two *T. aestivum* elite lines “Buster” (“Brimstone” x “Parade”; Nickerson Seeds Ltd.) and “Charger” (Fresco ‘Sib’ x Mandate; Plant Breeding International Cambridge Ltd.) using the standard wheat x maize technique from F1 plants (Laurie and Reymondie, 1991). These cultivars were chosen with the aim of obtaining a well-adapted population with regard to the UK climate, while segregating in flowering time. Both cultivars share common backgrounds and are semi dwarf (*Rht-D1b*) day length sensitive winter-type wheat varieties.

The Buster x Charger mapping population was grown as autumn (October) sown field experiments in the seasons 2011-2012, 2012-2013 and 2013-2014 across two locations (CF; John Innes Centre, Church Farm, Norwich, Norfolk, UK, and WP; Limagrain Ltd., Woolpit, Suffolk, UK), totalling in 4 unique year-sites (CF12, CF13, WP13, CF14). For the year-sites CF13, CF14 and WP13 the experiments were designed as randomized complete blocks with three replications and a plot size of 5.4m^2^ and 6m^2^ for the WP and CF experiments, respectively. In CF12, not enough seed was yet available for a replicated experiment. Thus, lines were drilled in two rows of 1m^2^ plots with random placement of genotypes. Crop husbandry was done according to standard agronomic practice in all experiments.

### Measurement of phenology

Exact determination of the terminal spikelet stage (TS) requires dissection of the shoot apex meristem and is often considered too laborious to be practical in larger field experiments. Thus, TS is usually determined approximatively, according to Zadok’s or BBCH growth stage 31, when the first node is detectable about 1cm above the tillering node. In this study, TS was nevertheless scored based on meristem dissections, to omit possible artifacts with regards to the exact timing of TS in the downstream analysis. In the CF12 experiment, every 2 – 3 days during tillering three random main tillers were selected from each plot and the apex development stage was scored through dissection in the field. A plot was considered to have reached TS if the terminal spikelet was observed in at least two of the three selected main tillers. Based on the observation in CF12, the sampling strategy was adjusted for the following CF13 and CF14 experiments. There, 5 random main tillers were collected for each plot on the same date (2013-05-09 and 2014-04-08, respectively), when the entire population was around TS. The segments containing the spikes for each plot were stored in FAA (water 50%, ethanol 35%, acetic acid 10%, and formaldehyde 5%, v/v; ; (Bancal, 2008)) for later dissection. TS stage was then recorded from the difference between the sampling date and the assigned TS date based on a back-scoring key developed from the CF12, CF13 and CF14 data (Fig. 1).

**Fig. 1:**
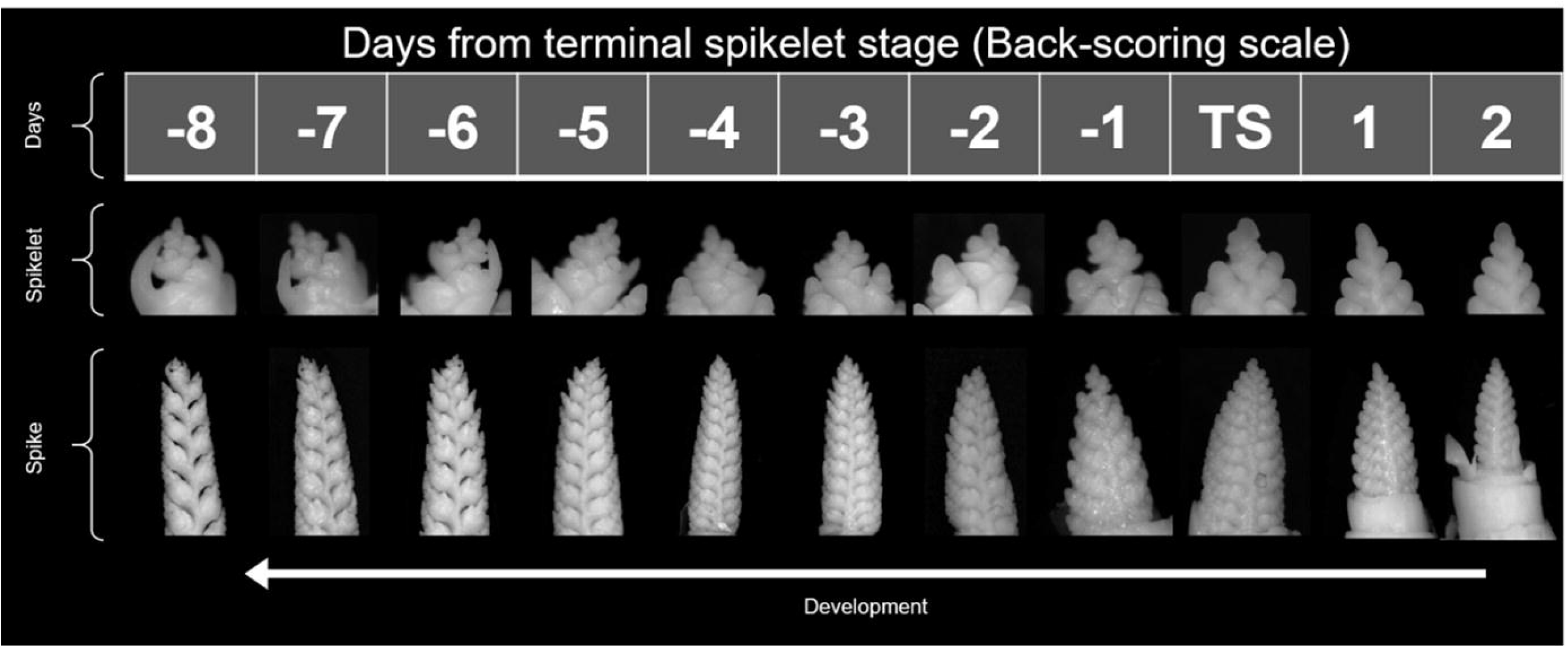
Back-scoring key for ex post determination of the terminal spikelet stage. Numbers denote the time difference in days from terminal spikelet stage (TS)

Ear emergence (EE) was recorded for each plot as days after sowing when at least 50% of spikes half emerged from the flag leaf sheath (BBCH 55;(Lancashire *et al*., 1991)). Stem elongation duration (SE) was recorded as the difference between EE and TS both in calendar days and in growing degree days, to account for the absolute duration of the phase as well as for putative effects of different temperature sums depending on the earliness of genotypes. Plant height was measured after all lines had reached final height by measuring three representative plants per plot from the base of the plant to the tip of the ear using a measuring stick.

### Measurement of yield components

At harvest, a sample of 10 spikes per plot were manually harvested. From these samples, number of grains per spike (G/spk), spikelets per spike (spklt/spk) and grains per spikelet (G/spklt) were manually counted and mean values across samples recorded as plot values. Grain yield for each plot was recorded by weighing the seed yield of the entire plot taken from the combine harvester and translating the values to t ha^-1^ based on the plot size. Thousand grain weight (TGW) was measured for each plot using a MARViN ProLine (MARViTECH GmbH, 19243 Wittenburg, Germany). Number of Grains per square meter (G/m^2^) was calculated from grain yield and TGW data for each plot. The number of spikes per square meter were calculated for each plot based on the G/spk and G/m^2^.

### Genotyping

Seeds of each line were germinated on water-soaked filter paper in a petri dish. One seedling per line was then planted in a plastic tray and moved to a greenhouse for 2 weeks when had developed 3-5 leaves. From each line, a 2cm leaf segment was cut and collected in a Qiagen 96 well box (Collection microtube, cat. No. 19560), placed on ice and stored at -80°C prior to extraction. The DNA was extracted following the protocol of Qiagen DNeasy 96 plant kit (cat. No. 69181). The extracted DNA was quantified using a Nanodrop spectrophotometer, measuring absorbance at 260nm and 280nm for quality (absorbance ratio) and concentration (compared to a standard curve), normalised to 50ng/ul and sent on dry ice to TraitGenetics (Am Schwabeplan 1b, Stadt Seeland OT Gatersleben, D-06466, Germany) for SNP genotyping and genetic map construction.

SNP genotyping was done using the illumina infinium 90K iSelect chip (Wang et al. 2014), resulting in a total of 5,665 SNP. The genetic map was generated using three different software packages (JoinMap 4.0, Map Manager QTXb20, and MapChart 2.2) and resulted in a total of 884 distinct loci across the genome, with an average distance of 5.7 cM between loci. The A and B sub-genomes had on average 54 and 50 loci per chromosome, respectively, whereas the D sub-genome had an average density of only 22 loci per chromosome (Supplementary Table S1). To facilitate downstream QTL analysis, the SNP haplotype blocks mapped to identical genetic loci were pooled and assigned a SNP group. Among the SNPs in a haplotype block, there was minimal allelic variation within individuals (occasional missing values or heterozygous alleles). Thus, for each individual, the prevalent allele among the SNPs in the group was assigned as SNP group allele. In case of ambiguity, the group allele was set to missing for the individual in question.

### Statistical Analysis

Statistical analysis was done in the R environment (R Core Team, 2024) following a two-stage approach (Kronenberg *et al*., 2021; Pérez-Valencia *et al*., 2022).

#### Single year-site analysis

In a first stage, phenotypic plot values for each trait and year-site were corrected for design factors and spatial trends using the R package SpATS (Rodríguez-Álvarez *et al*., 2018) parameterized with the model

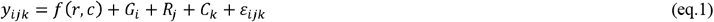

where *y*_*i,j,k*_ is the plot value of the *i*^th^ genotype in the *j*^th^ row and *k*^th^ column; *f*(*r, c*) is a smooth bivariate surface in the dimensions of rows and columns, which can be decomposed in a bilinear polynomial and a smooth part (PS-ANOVA, see Lee *et al*., 2013; Rodríguez-Álvarez *et al*., 2018); *G*_*i*_ is the fixed effect of the i^th^ genotype; *R*_*j*_ and *C*_*k*_ are the random effects of the *j*^th^ row and *k*^th^ columns, respectively and *ε*_*ijk*_ the plot residual.

Spatially corrected plot values were calculated as the genotypic best linear unbiased estimate (BLUEs) + plot residual (*ε*_*ijk*_) from eq.1. Within year-site repeatability was calculated according to Oakey et al. (2006), by setting Ci in eq.1 as random and applying ‘getHeritability’ of SpATS.

For the CF12 Experiment, eq.1 could not be fitted as there was only one replication and insufficient row/column arrangement in the experimental design. Thus, instead of spatially corrected values, the raw plot values of CF12 were taken for the second stage.

#### Across year-site analysis and heritability

In a second stage, across year-site best linear unbiased predictors (BLUPs) and heritabilities were estimated by fitting the model:

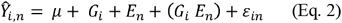

in the R-package Asreml-R (Buttler), where *Ŷ*_*i,n*_ are the spatially corrected plot values of the i^th^ genotype in the n^th^ year-site from the previous stage (eq.1), *µ* is a global intercept, *E*_*n*_ is the fixed effect of the n^th^ year-site *G*_*i*_ is the random genotype response, (*G*_*i*_ *E*_*n*_) is the random genotype by year-site interaction effect with a diagonal variance structure for the year-sites, allowing for year-site specific variances and *ε*_*in*_.is the residual variance. The variance structure of *G*_*i*_ was set according to identity by state (IBS) relationship structure among genotypes calculated from the SNP data using ‘snpgdsIBS’ function in the R-package SNPRelate (Zheng *et al*., 2012). To accommodate the covariance structure among genotypes, heritability was estimated based on genotype differences as 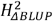 according to Eq. (24) in Schmidt et al. (2019).

#### Correlation analysis

Phenotypic correlations among across year-site BLUPs were calculated as Spearman’s rank correlation coefficient using the R-function ‘cor.test’. To calculate genetic correlations, Eq. 2 was expanded to the bivariate model:

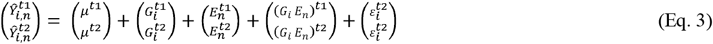

Where 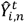 are the spatially corrected plot values for i^th^ genotype in the n^th^ year-site and the respective traits (t1 = trait 1, t2 = trait 2),*μ*^*t*1^ and *μ*^*t*2^ are the respective global intercepts and, 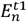 and 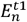 are the fixed year-site effects. 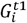 and 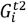 are the random genotype and (*G*_*i*_ *E*_*n*_)^*t*1^ and (*G*_*i*_ *E*_*n*_)^*t*2^ are the random genotype by year-site interaction effects with uniform variances per trait. Again, the variance structure of *G*_*i*_ was derived from IBS and uniform variances.

The estimated variance and covariance components from Eq. 3 were used to calculate the genetic correlations following Holland et al. (2001):

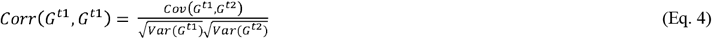

Where G denotes the genotypic variance (var) and covariance (cov) components of the respective traits (t1 = trait 1, t2 = trait 2)

#### QTL Mapping

In order to discern the genomic basis of the investigated traits, we applied QTL mapping implemented in the R package qtl (Broman *et al*., 2003) on the BLUEs of individual year-sites as well as the across year-site BLUPs. First, an initial genome scan was performed with a single QTL model using Haley-Knott regression by applying the function ‘scanone’ on imputed genotype data derived by applying the functions ‘calc.genoprob’ and ‘sim.geno’. Genotypes were imputed in 1cM intervals. Second, if putative QTL were detected, composite interval mapping was applied using the putative QTL from the first step as covariates using the function ‘cim’. For both the initial scans and composite interval mapping, permutation tests with 5000 permutations each were performed to establish the genome wide significance thresholds (α = 0.05) for every trait and year-site.

## Results

All investigated traits displayed moderate to high heritabilities across year-sites (0.37 ≤ H^2^ ≤ 0.95, Table 1) which was reflected in within year-site repeatabilities. Highest heritabilities were observed for phenology traits (TS, SE_d_, SE_GDD_, EE) and height (0.89 ≤ H^2^ ≤ 0.95) whereas lowest heritabilites were observed for grain and spike related yield traits (G/spk, G/spklt, spk/m^2^; 0.37 ≤ H^2^ ≤ 0.42). Yield, TGW, and G/m^2^ showed heritabilities of 0.66, 0.63 and 0.64, respectively.

**Table1.** Within year-site repeatability and across year-site heritabilities for all traits.

|  | CF12 | CF13 | WP13 | CF14 | all |
| --- | --- | --- | --- | --- | --- |
| trait | $H^2_{\text{Oakey}}$ | $H^2_{\text{Oakey}}$ | $H^2_{\text{Oakey}}$ | $H^2_{\text{Oakey}}$ | $H^2_{\Delta\text{BLUP}}$ |
| SE<sub>GDD</sub> | x* | 0.76 | -** | 0.86 | 0.92 |
| TGW | - | - | - | 0.57 | 0.63 |
| SE<sub>d</sub> | x | 0.71 | - | 0.81 | 0.89 |
| EE | x | 0.87 | - | 0.92 | 0.95 |
| TS | x | 0.8 | - | 0.83 | 0.93 |
| G/m<sup>2</sup> | - | 0.23 | - | 0.69 | 0.64 |
| G/spk | - | 0.7 | - | 0.64 | 0.4 |
| G/spklt | - | 0.15 | - | 0.63 | 0.42 |
| height | x | - | 0.93 | - | 0.91 |
| Spk/m<sup>2</sup> | - | 0.33 | - | 0.44 | 0.37 |
| spklt/spk | - | - | - | 0.86 | 0.88 |
| yield | - | 0.67 | 0.64 | 0.69 | 0.66 |

\* “x” denotes missing heritability due to impossible model fit
\*\* “-“ denotes missing heritability due to no trait data for the year-site

### Correlation analysis

To get an overview of the different traits and their interrelations, we conducted a correlation analysis among multi-environment BLUPs (Fig. 2).

**Figure 2:**
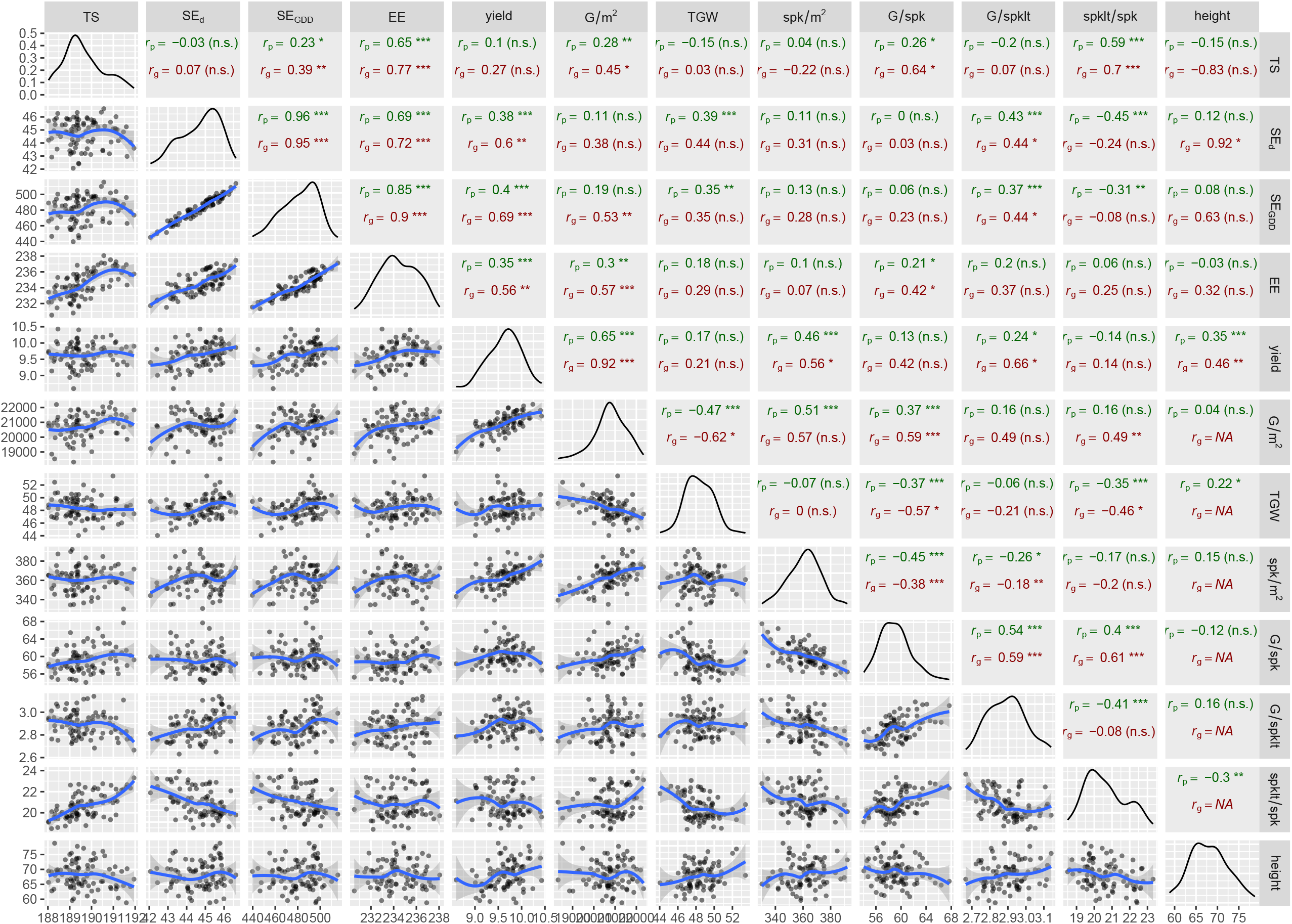
Correlation analysis among the investigated traits. The lower triangle displays scatterplots of BLUPs between respective trait pairs, including trend lines (blue) and standard errors (grey shaded areas) fitted using loess regression. The diagonal shows density distributions of the respective trait BLUPs. The upper triangle shows Spearman’s correlation coefficients (*r*_p_, green) as well as genetic correlation coefficients (*r*_g_, red) calculated from BLUPs of respective trait pairs. The significance of the respective correlation coefficients is indicated by: “***” = p < 0.001, “**” = p < 0.01 and “*” = p < 0.05, whereas “(n.s.)” denotes not statistically significant correlations.

### Phenology is mainly driven by heading time

The stem extension duration, irrespective of absolute or thermal time, was mainly driven by heading time (EE; *r*_g_ = 0.72, *r*_p_ 0.69 for SE_d_ and *r*_g_ = 0.9, *r*_p_ = 0.85 for SE_GDD_). Even though a strong correlation was found between EE and TS (*r*_g_ = 0.77, *r*_p_ = 0.65). The start of SE showed no significant correlation to SE duration in absolute time. However, a weak positive correlation was observed between TS and SE duration in thermal time (*r*_g_ = 0.39, *r*_p_ = 0.23), which highlights the usual, higher observed temperatures later in the season.

Plant height showed a strongly negative, albeit non-significant genetic correlation to TS (*r*_g_ = -0.83 (n.s.)) and a weak, non-significant positive correlation to EE (*r*_g_ = 0.32 (n.s.)), consistent with the very strong, significant correlation between height and SE_d_ (*r*_g_ = 0.92), and the strong but non-significant correlation between height and SE_GDD_ (*r*_g_ = 0.63 (n.s.)).

All phenotypic correlations between height and the phenology traits were weak (*r*_p_ ≤ 0.15) and non-significant. In summary, phenology appears to be largely driven by heading time and later lines have a tendency towards being taller and having longer stem extension duration.

### Grain number is more yield relevant than grain weight

Grain yield showed strong, positive genetic correlations to grains per square meter (G/m^2^; *r*_p_ = 0.65, *r*_g_ = 0.92), spikes per square meter (spk/m^2^; *r*_p_ = 0.46, *r*_g_ = 0.56) and plant height (*r*_p_ = 0.35, *r*_g_ = 0.46), but no significant correlations to TGW. The missing correlation between yield and TGW is in concordance with the fact that grain yield is often determined mainly by grain number rather than grain size under field conditions. Grains per square meter correlated strongly to spk/m^2^, however, the genetic correlation was non-significant (*r*_p_ = 0.51, *r*_g_ = 0.57 (n.s.)).

Among the yield components related to grain number, positive correlations were observed between G/spk on the one hand, and G/spklt (*r*_p_ = 0.54, *r*_g_ = 0.59) and spklt/spk (*r*_p_ = 0.4, *r*_g_ = 0.61) on the other. In accordance, G/m^2^ was positively correlated to G/spk (*r*_p_ = 0.37, *r*_g_ = 0.59) and genetically correlated to spklt/spk (*r*_p =_ 0.16 (n.s.), *r*_g_ = 0.49). However, G/spklt showed no significant correlation to G/m^2^, even though the genetic correlation was strong (*r*_g_ = 0.49 (n.s.)). Furthermore, G/spklt showed a negative phenotypic correlation towards spklt/spk (*r*_p_ = -0.41) but no genetic correlation was observed *r*_g_ = -0.08 (n.s.)). Similarly, spklt/spk and G/spk showed no direct correlation to grain yield (*r*_p_ = -0.14 (n.s), *r*_g_ = 0.14 (n.s) and *r*_p_ = 0.13 (n.s.), *r*_g_ = 0.42 (n.s.), respectively), whereas G/spklt did correlate to yield (*r*_p_ = 0.24, *r*_g_ = 0.66). While spk/m^2^ was positively related to total grain number, it showed negative correlations to grains per spike (*r*_p_ = -0.45, *r*_g_ = -0.38), grains per spikelet (*r*_p_ = -0.26, *r*_g_ = -0.18) and spikelets per spike, although the latter were non-significant (*r*_p_ = -0.17 (n.s), *r*_g_ = -0.2 (n.s.)). Therefore, increased spikes per unit area increase total grain number at the cost of grain number per individual spike. Similarly, more spikelets per spike lead to fewer grains per individual spikelet.

### Heading and SE duration drive yield

Stem extension duration in days (SE_d_; *r*_g_ = 0.6, *r*_p_ = 0.38) and SE duration in thermal time (SE_GDD_ r.g. = 0.69, *r*_p_ = 0.4) showed the strongest correlation to yield, followed by heading (*r*_p_ = 0.35, *r*_g_ = 0.56). Regarding the yield components, SE_d_ and SE_GDD_ duration were, phenotypically as well as genetically, positively correlated to G/spklt (0.37 ≤ r_*p,g*_ ≤0.44) and phenotypically negatively correlated to spklt/spk, while the respective genetic correlations were non-significant. Neither SE_d_ nor SE_GDD_ showed any significant correlations to G/spk or spk/m^2^. Yet, a significant genetic correlation was observed between SE_GDD_ and G/m^2^. Furthermore, both SE_GDD_ and SE_d_ showed positive phenotypic correlations to TGW (r_p_ = 0.35 and 0.39, respectively). The respective genetic correlations were of similar magnitude but non-significant.

Heading was positively related to G/m^2^ (*r*_p_ = 0.3, *r*_g_ = 0.57) and G/spk (*r*_p_ = 0.21, *r*_g_ = 0.42). The start of stem elongation showed positive relationships to G/m^2^ (*r*_p_ = 0.28, *r*_g_ = 0.45), G/spk (*r*_p_ = 0.26, *r*_g_ = 0.64) and spklt/spk (*r*_p_ = 0.59, *r*_g_ = 0.7). All other correlations between phenology and yield components were non-significant.

To further investigate the relationship between phenology and yield, we grouped genotypes by their yield quartiles (top 25%, central 50% and bottom 25%) and plotted yield versus start of SE, SE duration and heading (Figure 3). This revealed that the highest yielding genotypes had an optimal heading time of around 233-235 days after sowing, a longer SE duration (>43d) and a tendency towards earlier onset of SE. However, in contrast to SE duration and EE, the top yielding 25% of genotypes displayed the entire range of the population in terms of timing of TS.

**Figure 3:**
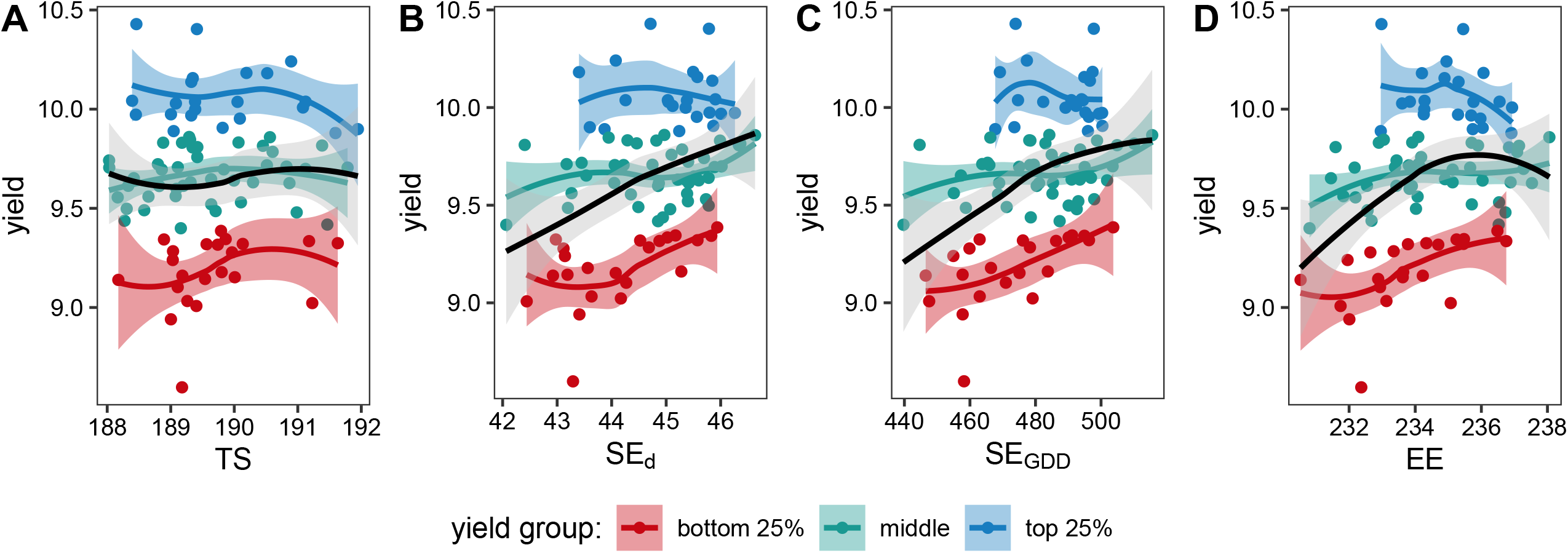
Relationship between phenology and yield among across year-site BLUPs. Genotypes were grouped according to their yield quartiles (blue = top 25 % of the population, green = middle 50 % of the population, red = bottom 25 % of the population). Red, green, and blue lines show loess trends for the respective yield groups, the black line shows the over-all trend among all data. Shaded areas depict respective standard errors of the loess trends. A-D: Scatterplots of yield versus start of stem elongation (A), stem elongation duration in days (B), stem elongation duration in thermal time (C) and ear emergence (D).

### QTL analysis

In total, QTL mapping detected 32 significant marker trait associations across chromosomes (chr) 1B, 1D, 2D, 3A, 4A, 4B and 7A for all investigated traits and year-sites except for the trait spk/m^2^ (Fig. 4, Supplementary table S2; LOD profiles, QTL effect profiles and QTL map locations are supplied in Supplementary File 1). For plant height, two independent QTL (i.e. not overlapping with any other QTL) were detected on chr 2D at 11cM (BS00065034_51) and on chr 4B at 1 cM (BS00009573_51), respectively. While the first was detected in the across year-site BLUPs as well as in the WP13 BLUEs for plant height, the latter was only detected in the CF12 experiment. Furthermore, one independent QTL each was detected for EE on chr 1B at 166 cM (snp_group_068) in the CF12 experiment, for SE_GDD_ on chr 1D at 74 cM (TA003135−0494) in the CF14 BLUEs, and for TGW on chr 4A at 0 cM (BS00035307_51) in the CF14 BLUEs.

**Fig 4:**
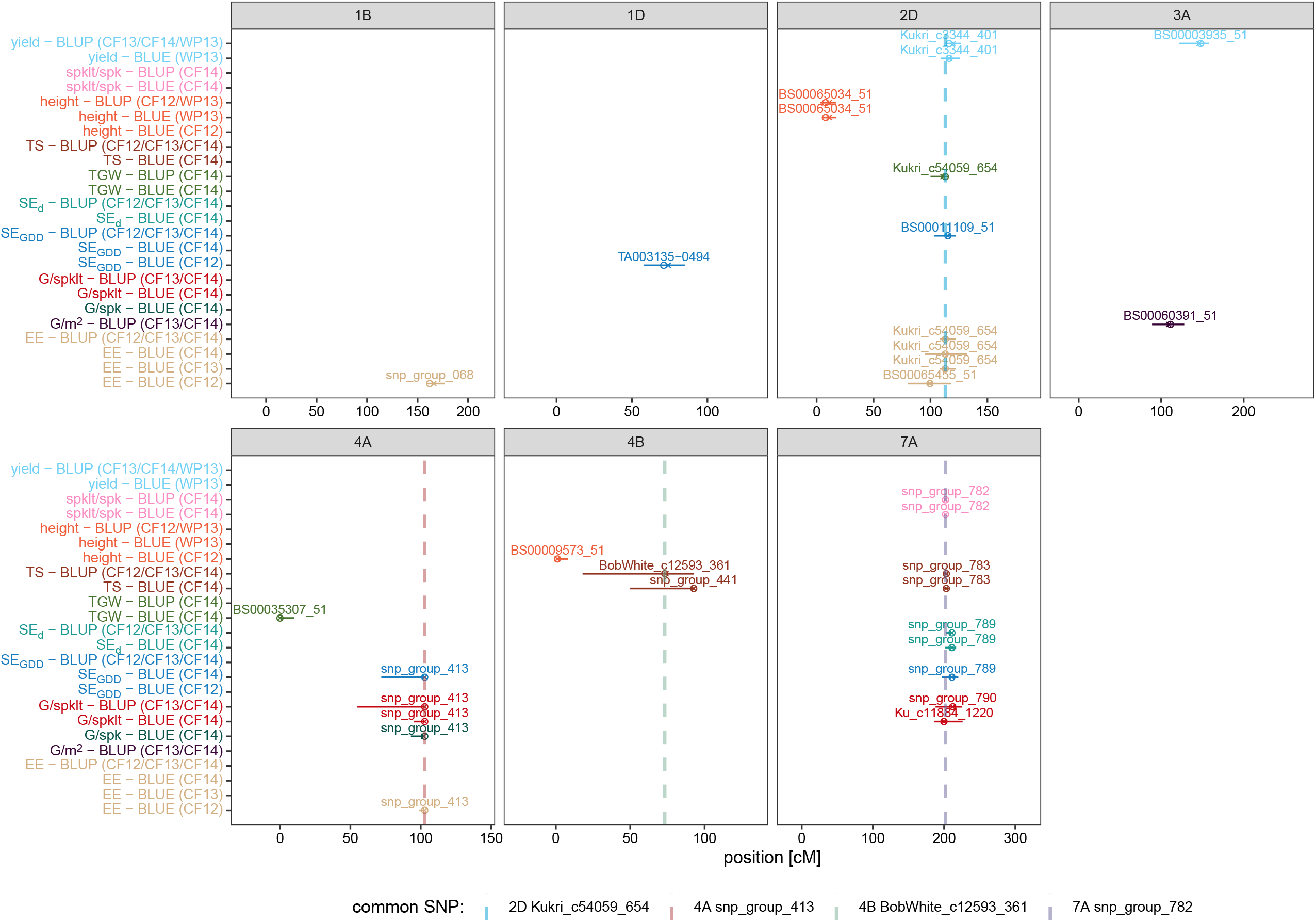
Detected marker-trait associations through QTL mapping. The x-axis shows chromosome locations in cM, grouped by chromosome (facets). The y-axis indicates trait – year/site pairs for the respective QTL. Refined QTL positions are given by x_s_ and positions of the nearest marker are given by open circles. Horizontal solid lines indicate the confidence intervals of the respective QTL. Colours of x_s,_ circles and horizontal lines indicate the trait of the respective QTL (see y-axis). Vertical dashed lines indicate the common SNP of overlapping QTL (see colour legend).

For TS, one QTL each was detected on chr 4B in the across year-site BLUPs (BobWhite_c12593_361 at 74 cM) and in the CF14 BLUEs (snp_group_441 at 92.7 cM). The confidence intervals of both associations largely overlapped (Fig. 4, Supplementary table S2). A yield QTL (across year-site BLUPs, BS00003935_51) together with a G/m^2^ QTL (across year-site BLUPs, BS00060391_51) was detected on chr 3A at 146 cM and 109 cM, respectively, whose confidence intervals showed a marginal overlap of 6 cM. Apart from these independent QTL, three clusters of mutli-trait QTL were detected on chr 2D, 4A and 7A, respectively (Fig. 4), reflecting the correlations observed in the data.

On chr 2D, the EE QTL Kukri_c54059_654 (CF13, CF14, across year-site BLUPs, overlapping with EE CF12 QTL BS00065455_51) also affected TGW (BLUPs CF14) and overlapped with the yield QTL Kukri_c3344_401 and the SE_GDD_ QTL BS00011109_51. On chr 4A, snp_group_413 was significantly associated with EE (BLUEs CF12), SE_GDD_ (BLUEs CF14), G/spk (BLUEs CF14) and G/spklt (BLUEs CF14, across year-site BLUPs). On chr 7A, the QTL snp_group_782 was associated with spklt/spk (CF14 BLUEs/BLUPs), and overlapped with QTL for TS (CF14, across year-site BLUPs), SE_d_ (CF14 BLUEs/BLUPs), GDD_SE_ (CF14 BLUEs) and G/spklt (CF14, across year-site BLUPs).

To gain more insight into the interrelations between phenology and yield traits and the respective individual and shared QTL for TS, EE and yield components, we looked at the respective allelic effects of snp_group_068 (EE; chr 1B), Kukri_c54059_654 (EE, SE_GDD_, TGW, yield; chr 2D), snp_group_413 (EE, SE_GDD_, G/spklt, G/spk; chr 4A), BobWhite_c12593_361 (TS; chr 4B) and snp_group_782 (TS, SE_d_, SE_GDD,_ G/spklt, spklt/spk; chr 7A) on the across year-site BLUPs of the respective traits (Fig. 5).

**Fig. 5.**
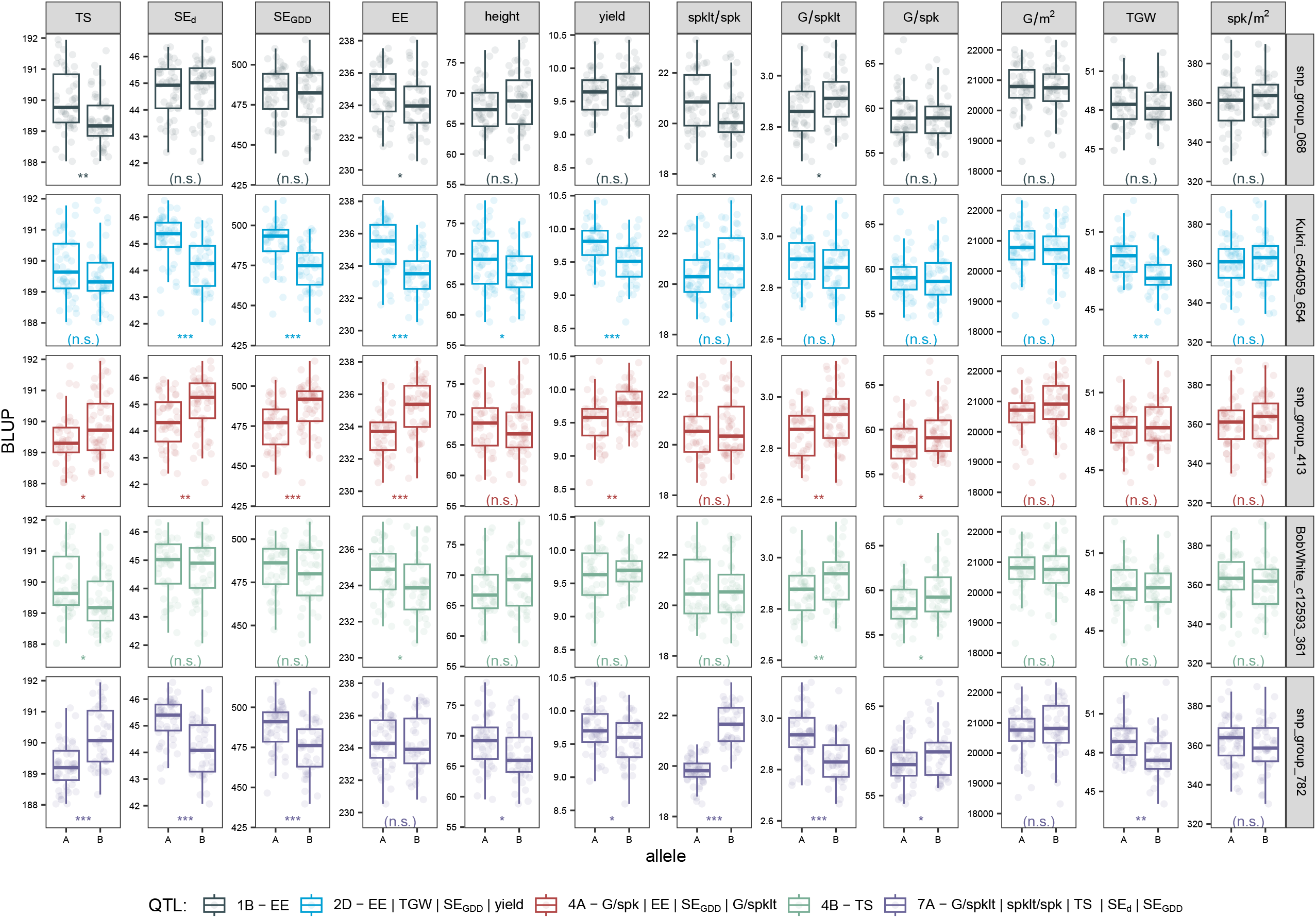
Effects of common SNP for overlapping QTL on the respective traits. Facet rows indicate the respective common SNP, coded by colour to indicate the associated traits (see colour legend). Boxplots and open circles show the BLUP values for the respective traits (facet columns) grouped by the marker alleles (A = Buster, B = Charger) The significance of the difference between groups calculated via T-test is indicated by: “***” = p < 0.001, “**” = p < 0.01 and “*” = p < 0.05, whereas “(n.s.)” denotes not statistically significant differences between groups.

Apart from its expected effect on EE, the 1B QTL snp_group_068 also significantly affected TS, spklt/spk and G/spklt. While both TS and EE were increased by the Buster allele, sklt/spk and G/spklt were affected in opposite directions, with the late Buster allele increasing spklt/spk but decreasing G/spklt. Consequently, no other effects on yield or yield components were registered. The 2D QTL Kukri_c54059_654 showed the expected allelic effect pattern on EE, SE duration, yield and TGW. While it also displayed a minor effect on plant height, it did not significantly affect TS or any of the grain number related yield components.

As expected, the 4A QTL snp_group_413 significantly affected EE, SE_GDD_, G/spklt and G/spk. In accordance with the observed correlations, TS, SE_d_ and yield also showed significant differences between the Buster and Charger alleles. The allelic effects were consistent with the expected pattern that increased SE duration is associated with more grains per spike. However, this did not translate into a significant effect on G/m^2^.

Apart from the expected effect on TS, the 4B QTL BobWhite_c12593_361 also affected EE, G/spk and G/spklt. In contrast to the chr 4A QTL but in accordance with the 1B QTL, here, the earlier allele led to more G/spk and G/spklt. Nevertheless, no further effects on yield or yield components were registered.

The 7A QTL snp_group_782 was the only detected locus that had a significant effect on TS without affecting EE. Furthermore, lines with the early allele for TS had significantly longer SE duration, were significantly taller and had significantly higher yield. However, the yield effect appears to be due to significantly higher TGW rather than higher G/spklt in the earlier lines compared to the late lines, as G/spk was lower in the early lines, due to a lower number of spklt/spk.

In summary, the 1B QTL and the 4B QTL affected TS and EE without affecting SE duration or yield. The 2D QTL affected EE and yield through TGW. The 4A QTL affected yield through grain number and SE duration traits. The 7A QTL affected TS and SE duration without affecting EE and lead to higher yield in the early TS lines due to higher TGW but not due to higher grain number, as number of spikelets per spike was drastically lower in the lines carrying the early allele.

## Discussion

In this study we measured the timing of SE and its relationship to grain yield and yield components in a sizeable wheat mapping population under field conditions across multiple years and locations, to determine whether and how a prolonged SE duration translates to increased yield. The selected population originated from a cross between two modern elite wheat cultivars well adapted to the UK climate. Thus, we consider the findings of this study relevant and transferable to current elite germplasm, at least for UK climatic conditions.

### High heritabilities indicate potential for selection

The presented back-scoring method allowed the accurate determination of the terminal spikelet stage in 108 genotypes across three year-sites, as evidenced by the high within year-site repeatabilities and the high across year-site heritability. In general, the observed heritabilities where within the expected ranges even though the heritabilities for G/m^2^, G/spk G/spklt and spk/m^2^ were only based on two individual year-sites and heritabilities for TGW and spklts/spk were based on one single year-site, respectively.

### Longer stem elongation duration increases yield through grain number

Investigating the phenotypic relationship between grain yield and the phenology traits clearly demonstrated the importance of an appropriate heading date, which is a well-known, major selection criterion in breeding for a specific environments (Jung and Müller, 2009). However, the results also showed a clear positive relationship between the duration of stem elongation and yield, based on phenotypic as well as genetic correlations. Furthermore, both yield and SE duration were positively genetically linked to G/m^2^. The link between SE duration and G/m^2^ was further supported by positive correlations to G/spklt. This results aligns well with previous findings, that grain number in response to assimilate competition is most affected by the number of spikelets per spike (Slafer *et al*., 1994; Backhaus *et al*., 2023; Anderegg *et al*., 2024). Together, these results provide the first field-based evidence from a substantial number of genotypes that extending SE duration enhances wheat yield through increased G/m^2^, validating a widely proposed physiological breeding strategy (Slafer and Rawson, 1994; Slafer *et al*., 1996, 2001*b*, 2015; Miralles and Slafer, 2007; Whitechurch *et al*., 2007; Reynolds *et al*., 2009; Pérez-Gianmarco *et al*., 2018). Furthermore, the results are also in concordance with the often reported negative relationship between grain number and grain weight, and the importance for grain number as a yield determinant under field conditions (Magrin *et al*., 1993; Slafer *et al*., 2001*a*; Peltonen-Sainio *et al*., 2007; Quintero *et al*., 2018).

### A link between stem elongation duration and grain weight

Apart from the relationship between SE duration and grain number, increased SE duration was also associated with higher TGW, as reflected in the phenotypic correlations and the effects of the 2D QTL Kukri_c54059_654 and the 7A QTL snp_group_782. Grain number and grain weight are often regarded as trade-off traits, with the former being set before anthesis and the latter being set during grain filling. However, increasing evidence suggest that the critical periods for grain number and grain weight overlap, as potential grain weight is established already between booting and grain setting, when carpels develop (Calderini *et al*., 1999, 2021; Vicentin *et al*., 2024). It was shown that increased temperatures during this period reduce grain weight (Calderini *et al*., 1999; Ugarte *et al*., 2007). A recent study found that grain weight was affected by temperature and global radiation from as early as floral transition (Sabir *et al*., 2023). We are not aware of any studies that previously associated TGW with SE duration. However, Xie and Sparkes (2021) found that earlier anthesis lead to increased TGW in Forno x Oberkulmer RIL population (Messmer *et al*., 1999). In contrast, our results suggest that an extended SE duration, either through later heading or through earlier TS, increased TGW. Importantly, this did not affect grain number and thus increased yield.

### Heading date is a major driver of phenology

The observed strong relationship of EE towards the start of SE and SE duration as well as the relationship between the phenology traits and height is in agreement with previous studies in European elite germplasm of more diverse origin (Kronenberg *et al*., 2017, 2021; Roth *et al*., 2023*b,a*). This, together with the weak relationship between the start of SE and SE duration indicates that phenology is strongly driven by heading time.

In terms of selecting for an increased SE duration to increase yield, this indicates a possible impasse. Delaying heading time would shorten the grain filling period and thus further increase the trade-off between grain number and grain weight and furthermore risk yield loss due to terminal drought, depending on the environment (Dorrani-Nejad *et al*., 2022; Zheng *et al*., 2022). On the other hand, selecting for later heading might be useful in environments with low risk of heat or terminal drought stress and counteract the tendency towards earlier heading due to global warming (Wang *et al*., 2015). Yet, the results of our QTL analysis indicate possibilities to fine-tune the stem elongation phase and yield independently.

### Independent QTL indicate possibilities to adjust TS without affecting heading

The detected independent and shared QTL among the investigated traits largely reflect the pattern observed in the correlation analysis. However, investigating their allelic effects including ‘off-target’ effects on non-associated traits indicates additional leeway. Fine-tuning of phenology and its effects on yield components may be possible through specific combinations of respective QTL alleles.

The 4A QTL (snp_group_413) is representative of the broader correlation pattern. Selecting for the Charger allele would lead to later varieties (both in TS and EE) with a longer SE phase and a higher yield due to increased grain number. In contrast, selecting for the Buster allele in the 7A (QTL snp_group_782) would lead to varieties with a longer SE duration due to earlier TS, without affecting EE or grain number but higher yield due to larger TGW. A selection of the Buster allele in the 2D QTL (2D Kukri_c54059_654) would lead to a longer SE duration due to later EE and increase yield without affecting grain number through increased TGW. The 7A and 2D QTL would also result in taller varieties whereas height is unaffected in the 4A QTL. Finally, selecting the respective Buster alleles in the 1B (snp_group_068) or the 4B (QTL BobWhite_c12593_361) QTL would lead to earlier varieties (both in TS and EE) without affecting grain number or grain weight.

## Conclusion and outlook

We set out to investigate whether a longer stem elongation indeed leads to increased yield through increased grain number and whether stem elongation duration can me modified by independently adjusting the start of stem elongation and heading. The results show that a longer stem elongation duration indeed increases yield through increased grain number. Furthermore, we found a positive relationship between stem elongation duration and grain weight, increasing over all yield. We present QTL that allow to independently finetune the timing of phenology and its effects on grain number and grain weight. Although these QTL need to be tested and confirmed in broader genetic backgrounds, they may offer the opportunity for further yield increase in a physiological breeding framework.

## Supplementary data

Table S1: Genetic map summary table

Table S2: Summary table of detected QTL for all traits and year-sites.

Supplementary File 1: QTL mapping results figures depicting LOD profiles, refined QTL profiles, estimated marker effects and map locations for all traits and year-sites.

## Acknowledgements

We would like to thank Lesley Fish and John Snape for developing and providing the double haploid population. Further, we thank Simon Berry for supporting the field trial at Woolpit.

## Author Contributions

**LK:** Methodology, Software, Formal analysis, Visualization, Funding acquisition, Writing - Original Draft.

**OG:** Investigation, Data Curation, Methodology, Writing - Review & Editing.

**SC:** Investigation.

**MC**: Investigation.

**PT:** Investigation.

**ML:** Investigation.

**LW:** Software, Methodology, Writing - Review & Editing

**SG:** Conceptualization Resources, Supervision, Project administration, Funding acquisition, Writing - Review & Editing.

All authors have read and approved the final manuscript.

## Data availability

All data are available from the authors upon reasonable request.

